# Dopaminergic modulation of dynamic emotion perception

**DOI:** 10.1101/2022.03.02.482469

**Authors:** B.A. Schuster, S. Sowden, A.J. Rybicki, D.S. Fraser, C. Press, P. Holland, J.L. Cook

## Abstract

Emotion recognition abilities are fundamental to our everyday social interaction. A large number of clinical populations show impairments in this domain, with emotion recognition atypicalities being particularly prevalent among disorders exhibiting a dopamine system disruption (e.g., Parkinson’s disease). Although this suggests a role for dopamine in emotion recognition, studies employing dopamine manipulation in healthy volunteers have exhibited mixed neural findings and no behavioural modulation. Interestingly, whilst a dependence of dopaminergic drug effects on individual baseline dopamine function has been well established in other cognitive domains, the emotion recognition literature so far has failed to account for these possible interindividual differences. The present within-subjects study therefore tested the effects of the dopamine D2 antagonist haloperidol on emotion recognition from dynamic, whole-body stimuli while accounting for interindividual differences in baseline dopamine. 33 healthy male and female adults rated emotional point-light walkers (PLWs) once after ingestion of 2.5 mg haloperidol and once after placebo. To evaluate potential mechanistic pathways of the dopaminergic modulation of emotion recognition, participants also performed motoric and counting-based indices of temporal processing. Confirming our hypotheses, effects of haloperidol on emotion recognition depended on baseline dopamine function, where individuals with low baseline dopamine showed enhanced, and those with high baseline dopamine decreased emotion recognition. Drug effects on emotion recognition were related to drug effects on movement-based and explicit timing mechanisms, indicating possible mediating effects of temporal processing. Results highlight the need for future studies to account for baseline dopamine and suggest putative mechanisms underlying the dopaminergic modulation of emotion recognition.

**Significance statement:** A high prevalence of emotion recognition difficulties amongst clinical conditions where the dopamine system is affected suggests an involvement of dopamine in emotion recognition processes. However, previous psychopharmacological studies seeking to confirm this role in healthy volunteers thus far have failed to establish whether dopamine affects emotion recognition and lack mechanistic insights. The present study uncovered effects of dopamine on emotion recognition in healthy individuals by controlling for interindividual differences in baseline dopamine function and investigated potential mechanistic pathways via which dopamine may modulate emotion recognition. Our findings suggest that dopamine may influence emotion recognition via its effects on temporal processing, providing new directions for future research on typical and atypical emotion recognition.

## Introduction

The ability to recognize others’ emotions from facial and bodily cues is an important skill that facilitates the development of meaningful social relationships (Izard et al., 2001; Wang et al., 2019). However, the cues towards genuine expressions of emotions are often subtle. When we are sad we do not pull a face that bears much resemblance to the Ekman example, but rather subtly alter the particular spatiotemporal dynamics of the way we move our body and face (e.g., Michalak et al. (2009); Roether et al. (2009); Sowden et al. (2021)). Therefore, it is perhaps unsurprising that difficulties labelling emotions are widespread throughout a wide range of clinical conditions, but most notably, within those featuring a disruption of the dopaminergic neurotransmitter system (e.g., Parkinson’s disease: Argaud et al. (2018), Huntington’s disease: Henley et al. (2012), schizophrenia: Edwards et al. (2002)). These observations warrant a closer examination of the role played by dopamine in socio-cognitive skills such as emotion recognition.

An incisive way to establish a causal role is to observe the influence of dopaminergic drugs on emotion recognition in the healthy population. Interestingly, while studies have found a range of mixed influences on neural responses (e.g., amygdala activation; Hariri (2002); Takahashi et al. (2005); Delaveau et al. (2007)) during emotion processing, they have typically not found behavioral influences. One explanation for this mixed picture is that there is an optimal level of dopamine for such tasks, and that – dependent upon one’s baseline levels – dopaminergic modulation brings individuals closer to, or further away from, that optimum (Delaveau et al., 2005; Delaveau et al., 2007). This theory has received widespread attention in other domains of cognition (Kimberg et al., 1997; Mattay et al., 2000; Gibbs and D’Esposito, 2005; Frank and O’Reilly, 2006; Cools and D’Esposito, 2011). Direct examinations of dopamine synthesis capacity and/or receptor binding are possible via positron emission tomography (PET) but are expensive and difficult to implement. Consequently, a large number of cognitive studies approximate striatal dopamine synthesis capacity via measures of working memory span – which are found to comprise a good proxy (Cools et al., 2008; Landau et al., 2009). Such studies find that individuals with low working memory span – signaling low dopamine synthesis capacity – exhibit different behavioral responses to a dopaminergic modulation to those with higher span (capacity).

It is also notable that while a role for dopamine in emotion perception is suggested by studies illustrating aberrations in clinical conditions with dopamine dysfunctions, we do not have good mechanistic models of the nature of that role. Some plausible contenders could relate to the influence of dopamine on temporal and motor encoding. Many of the cues signaling emotional state are dynamic, and will therefore depend upon one’s ability to encode temporal features. For instance, whereas rapid, accelerated movements are associated with anger, slower and sluggish movements tend to be interpreted as sadness (Gross et al. (2012); Edey et al. (2017); Halovic and Kroos (2018)). It has also been hypothesized – outside of the dopamine literature – that recognition of such temporal features relies upon yoking to one’s own movements and the emotional state experienced when performing such movements (Edey et al., 2017; Edey et al., 2020). Given the strong link between dopamine and temporal encoding (Coull et al., 2012; Tomassini et al., 2016), as well as motor performance (Niv et al., 2007; Tomassini et al., 2016), it is plausible that dopamine mediates emotion recognition via its influence on temporal processing.

To examine the role played by dopamine in emotion recognition, this study presented participants with a dynamic whole-body emotion recognition task under the dopamine D2 receptor antagonist haloperidol, and a placebo. We separated our analyses according to baseline working memory span and examined whether influences of the drug were modulated by performance in movement- and counting-based indices of temporal processing.

## Method

### Participants

Forty-three healthy volunteers (19 females; mean (M) [SD] age = 26.36 [6.3]) took part on at least one of two study days after passing an initial health screening. Participants were recruited via convenience sampling from University of Birmingham campus and Birmingham city center, gave written informed consent and received either money (£10 per hour) or course credit for participation. Five participants (2 placebo, 3 haloperidol) dropped out of the study after completing the first day, a further five could not complete the second test day due to COVID-19 related closures, consequently all analyses are based on 33 full datasets. All experimental procedures were approved by the University of Birmingham Research Ethics Committee (ERN 18-1588) and performed in accordance with the World Medical Association Declaration of Helsinki (1975).

### Experimental design and statistical analyses

#### Pharmacological manipulation and general procedure

Participants’ eligibility for the study was evaluated by a clinician via review of their medical history, electrocardiogram assessment and blood-pressure check. The main study took place on two separate test days, one to four weeks apart, where participants first completed an initial blood-pressure and blood-oxygenation check with the medic. Subsequently, in a double-blind, placebo controlled within-subjects design, each participant took part on two study days, wherein all participants received tablets containing either 2.5 mg haloperidol or lactose (placebo) on the first day, and the respective other treatment on the second day (order of drug day counterbalanced). For this, participants were handed a pre-prepared envelope, instructed that this would contain either placebo or haloperidol tablets, informed that none of the experimenters knew the contents of the envelope, and asked to close their eyes before swallowing the tablets. Haloperidol is a dopamine D2 receptor antagonist, which affects dopamine transmission via binding either to postsynaptic D2 receptors (blocking the effects of phasic dopamine bursts), or to pre-synaptic autoreceptors (which has downstream effects on the release and reuptake of dopamine and thus modulates bursting itself; Benoit-Marand et al. (2001); Schmitz et al. (2003)). Effects of dopaminergic agents can vary depending upon an individual’s baseline dopamine synthesis capacity, potentially due to increased drug sensitivity in those with low synthesis capacity resulting from upregulated receptor density and/or sensitivity (e.g., Cools et al., 2008; Landau et al., 2009).

Reported mean values for peak concentration and elimination half-life of oral haloperidol lie between 1.7 and 6.1 and 14.5 – 36.7 hours, respectively (Kudo and Ishizaki, 1999). After drug administration, participants rested for 1.5 hours to allow for drug metabolization. Subsequently participants began the task battery, which included the emotion recognition task, a verbal working memory task and indices of drug effects on movement- and counting-based temporal processing (see Method: Tasks and procedure). Throughout the day, participants’ blood -pressure and -oxygenation was checked hourly between tasks. All data was collected at the Centre for Human Brain Health (CHBH) at the University of Birmingham, UK.

#### Tasks and procedure

Participants completed a task battery including tasks not described in this study. All relevant tasks are described below in order of presentation. Task order was the same on both study days.

##### Visual working memory (WM) task

This task was implemented to establish a proxy for baseline dopamine synthesis capacity. Specifically, PET studies have shown that low working memory scores are associated with low dopamine synthesis capacity, and thus have been suggested to reflect higher susceptibility to the effects of dopaminergic drugs (Cools et al., 2008). Correspondingly, for many cognitive tasks, behavioral effects of dopaminergic drugs are found to be different in individuals with low and high baseline working memory span (typically determined based on median-split: Kimberg et al. (1997); Mattay et al. (2000); Gibbs and D’Esposito (2005); Frank and O’Reilly (2006)). For example, on a number of tests of executive function, performance is boosted by dopaminergic drugs in low-span participants, and impaired in high-span participants (Kimberg et al., 1997; Mattay et al., 2000; Gibbs and D’Esposito, 2005).

Participants completed an adaptation of the Sternberg (Sternberg, 1966) visual WM task. Participants completed 60 trials across five blocks. On each trial, a fixation cross was displayed at the center of the screen (500-1000 ms), followed by a list of consonants (5 – 9 characters in length, depending on the block; 1000 ms), followed by a blue fixation cross (3000 ms). A single test letter was then displayed (4000 ms), and participants were asked whether the letter was taken from the previously displayed list (Fig. 1B). Participants responded by pressing 1-3 on the keyboard (1 – Yes, 2 – No, 3 – Unsure). Responses (accuracy) and response time (time from test letter displayed until participant response) were recorded for each trial. Each block varied in length from 5-9 consonants, with letters randomly selected from the alphabet on each trial. The total task duration was approximately 10 minutes and test trials were preceded by 10 practice trials.

**Figure 1.**
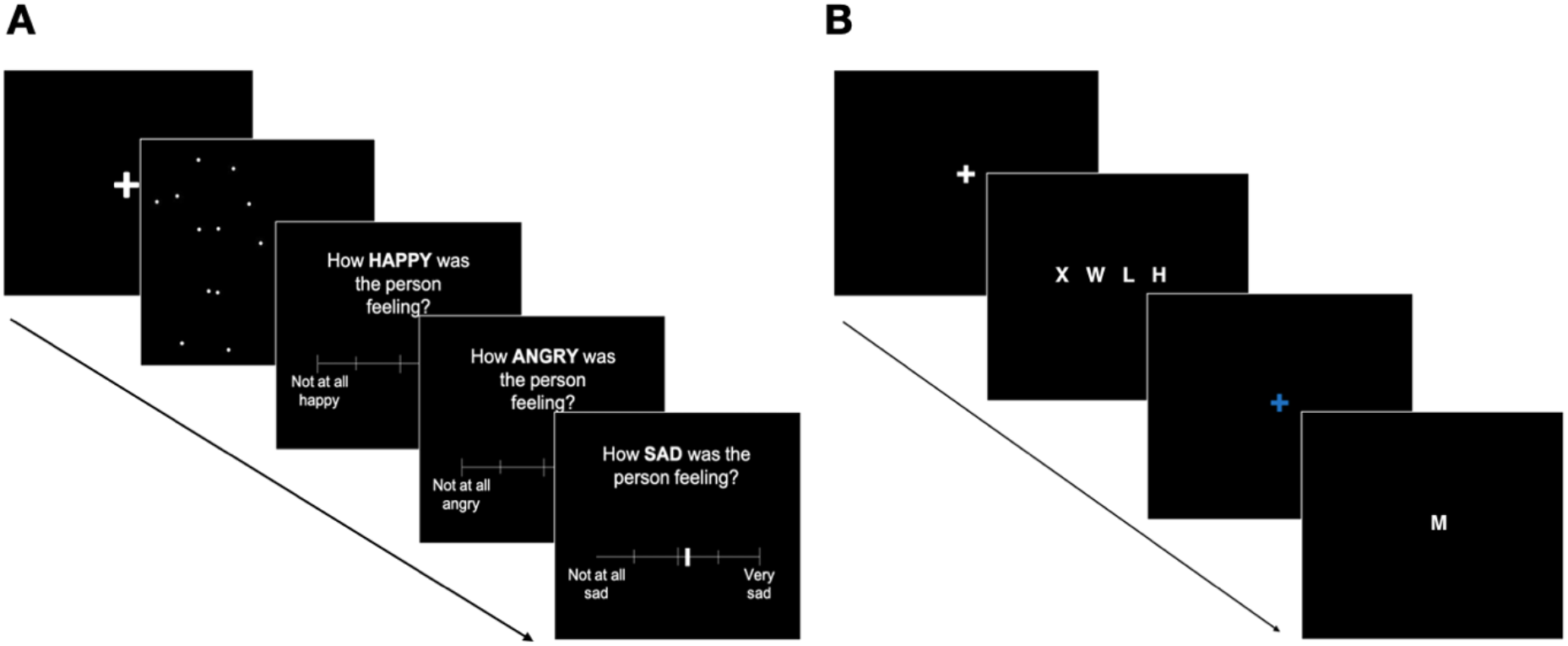
Schematic depiction of main tasks. (A) PLW perception task. (B). Visual working memory task. *Note.* (A) Depiction of one trial of PLW perception task. A fixation cross was presented for 1000 ms and followed by a PLW stimulus (on average 2000 ms). Subsequently participants rated on three separately presented scales (each ranging from ‘Not at all’ to ‘Very’) in pseudorandom order how angry, happy, and sad they perceived the PLW stimulus to be. (B) Depiction of one trial of visual working memory task. After presentation of a fixation cross (duration varied between 500-1000 ms), a list of 5-9 characters was presented for 1000 ms, followed by a blue fixation cross (3000 ms).

##### Time estimation task

In the time estimation task, participants were asked to estimate temporal intervals of varying lengths by counting the number of seconds that had passed between two auditory signals. Four time intervals of varying lengths (between 22 and 103 seconds) were presented in a pseudorandom order.

##### Emotion recognition task

Stimuli were whole-body point light displays of male and female actors modelling angry, happy and sad emotional walks (i.e., point light walkers [PLWs]) adopted from Edey et al., (2017)). For each of the three affective states, the stimulus set contained 100% stimuli, which displayed the walkers at the speed they originally modelled. In line with the literature demonstrating that sadness is conveyed via slow, sluggish movements, anger with fast, jerky kinematics, and happiness intermediate to the two (Michalak et al., 2009; Roether et al., 2009; Gross et al., 2012; Halovic and Kroos, 2018), sad 100% PLWs exhibited the slowest mean speed, followed by happy, and then angry PLWs (Nackaerts et al., 2012). In addition, for each emotion, the stimulus set included three levels of velocity adapted stimuli, consisting of morphs between the speed of neutral walkers and the corresponding 100% stimuli. The resulting velocity adapted stimuli thus contained 0%, 33% and 67% of emotion specific speed information with full postural information. A total of 48 velocity adapted and 100% stimuli (4 trials of angry, happy, and sad PLWs at 4 levels of speed information) were presented in pseudorandom order for an average of 2000 milliseconds (ms). On each trial, participants first viewed a fixation cross for 1000 ms, followed by a PLW stimulus. Subsequently three separate visual-analogue scales (ranging from 1 [not at all] to 10 [very]) were presented one after another, in pseudorandom order, asking participants to rate how intensely they felt the stimulus was expressing an angry, happy, or sad emotional state (Fig. 1A).

##### Walking task

Following the emotion recognition task (task order was fixed to avoid priming effects of own speed on emotion judgements from PLWs’ speeds; for more details see Edey et al. (2017)), participants performed the walking task. Individuals were asked to walk continuously between two sets of cones (placed 10 meters apart) for 120 seconds at their preferred walking speed. Each participant completed two walks (of 120 seconds) approximately 30 minutes apart. Acceleration data was recorded, using the SensorLog app^1^, with an iPhone 6s attached to the outer side of the participants’ left ankle.

#### Statistical analyses

All data were processed in Matlab R2021a (The MathWorks Inc., 2021) and analyzed with Bayesian linear mixed effects models using the *brms* package (Bürkner, 2017) in R (version 4.2, R Core Team, 20201). Prior to model building, any continuous predictors were normalized to allow comparisons between individual estimates. For all analyses, model building was guided by our experimental design and the final model selected based on leave-one-out cross validation (using the ‘LOO’ subfunction of brms; for details on all models compared see ‘Code and data accessibility’). The contribution of individual predictors to the model was evaluated based on the posterior probabilities of their expected values and confirmed by comparing the model to one excluding the predictor in question. For all relevant model parameters we report expected values under the posterior distribution and their 95% credible intervals (CrIs^2^), as well as the posterior probability that a certain effect (*Eµ*) is different from zero (P(Eμ < 0) / P(Eμ > 0)). In line with Franke & Roettger (2019), if a hypothesis states that an effect is not equal to zero (*Eµ* ≠ 0), we conclude there is compelling evidence for this effect if zero is not included in the 95% CrI of *Eµ* and if the posterior probability P(*Eµ* ≠ 0) is close to 1. We used weakly informative priors, following a normal distribution for the intercept and all regression coefficients and a half-Cauchy distribution for residual and random effect variances (all prior distributions were centered at 0 and had a standard deviation of 10; see Nalborczyk et al. (2019)). For all models, the maximal possible random effects structure allowed by the design was defined (Barr et al., 2013). Each model was run for four sampling chains with 5000 iterations each (1000 warm-up iterations). There were no indications of nonconvergence (all Rhat values = 1, no divergent transitions).

#### Code and data accessibility

All data and code required to reproduce the analyses within this article is publicly available under https://osf.io/gcvyj/.

## Results

Based on previous evidence indicating that WM span reliably predicts individual dopamine synthesis capacity (Cools et al., 2008; Landau et al., 2009) and drug effects on performance in other cognitive domains (Kimberg et al., 1997; Mattay et al., 2000; Gibbs and D’Esposito, 2005; Frank and O’Reilly, 2006; Rostami Kandroodi et al., 2021), we used individual *baseline WM span* as a proxy for baseline dopamine function and thus interindividual differences in drug responsivity. For this, WM task accuracy was calculated as the percentage of correct responses of all placebo trials. Groups of low and high baseline WM span (indexing low and high dopamine synthesis capacity, respectively) were then defined by performing a median split on WM span scores from the placebo day only (resulting in 18 and 20 subjects in the low and high WM groups, respectively).

### Emotion recognition task

As in Edey et al. ((Edey et al., 2017)), *emotion recognition scores* were calculated for each emotion and speed level by subtracting the mean of the ratings for the two non-modelled emotions from the rating for the modelled emotion. For example, for a sad PLW stimulus, we subtracted the mean of the ratings on the angry and happy scales from the rating given on the sad scale. Possible values thereby ranged from -9 to 9, with high emotion recognition scores reflecting judgements of the PLW intensely expressing the modelled emotion and successful discrimination between the three emotion scales, and low or negative emotion recognition scores indicating that participants felt the PLW was weakly expressing the modelled emotion or a lack of discrimination between the three emotion scales.

#### Haloperidol increased emotion recognition scores in low WM span, and decreased emotion recognition in high WM span individuals

To confirm that individuals made use of the emotion-specific speed information when rating PLW stimuli, and to ascertain that the drug did not affect particiants’ sensitivity to the speed manipulation, an initial control model was conducted: A Bayesian linear mixed effects model with a random intercept for *subject ID* was fitted to the factors *drug* (placebo [PLA], haloperidol [HAL]; dummy coded), *emotion* (sad, happy, angry; effects coded), *speed level* (i.e., emotion specific speed information; 0%, 33%, 67%, 100%; orthogonal polynomial coded) and *WM group* (low, high; effects coded), as well as all possible two- and three-way interactions, predicting emotion recognition scores. Due to the dummy-coding of the factor drug all main effects refer to the placebo level, which are compared to effects under haloperidol via individual contrasts. The control model revealed a strong positive linear trend for the variable speed level (*Eµ*_*PLA,speedLevel*.*L*_ = 1.29, CrI = [0.94, 1.64]), confirming that, overall, participants gave increasing emotional intensity ratings with increasing speed levels. There were no interactions between drug and speed level, indicating that Haloperidol did not affect participants’ sensitivity to the speed manipulation. Consequently, all following results are reported based on emotion recognition scores collapsed across the four speed levels.

In the subsequent model (Bayesian linear mixed effects model with random intercept for subject ID, factors drug, emotion and WM group, dependent variable emotion recognition scores collapsed across speed level), there was a main effect of emotion for PLA trials, with contrasts revealing that overall, sad PLWs were rated with higher intensity (*Eµ*_*PLA,sad*_ = 0.65, CrI = [0.36, 0.93]), while angry PLWs were rated lower than average in terms of intensity (*Eµ*_*PLA,angry*_ = -0.59, CrI = -0.88, -0.31). There was no main effect of drug, with the contrast of PLA and HAL emotion recognition scores being close to zero (*Eµ*_*PLA-HAL*_ = -0.06 (CrI = [-0.37, 0.24]).

Most interestingly and confirming our primary hypothesis, there was an interaction between drug and WM group, with a 0.94 point difference between drug effects on emotion recognition scores in the low and high WM group (*Eµ*_(*PLA-HAL,lowWM*)-(*PLA-HAL,higfiWM*)_ = - 0.94, CrI = [-1.56, -0.32]). To evaluate drug effects in the two WM groups, two separate post-hoc models were run for low and high WM groups. These models confirmed the predicted nature of differences, revealing superior performance under haloperidol versus placebo in the low WM group (*Eµ*_*PLA-HAL,lowWM*_ = 0.40, CrI = [-0.06, 0.87]), probability that this is a true effect: P(*Eµ*_*PLA-HAL,JhighWM*_ > 0) = 0.96), alongside poorer performance under haloperidol in the high WM group (*Eµ*_*PLA-HAL,highWM*_ = -0.53, CrI = [-0.95, -0.12]), probability that this is a true effect: P(*Eµ*_*PLA-HAL,highWM*_ < 0) = 0.99; see Fig 2). These improvements under haloperidol in the low WM group were generated via increased ratings on the modelled scales and decreased ratings on the non-modelled scales – demonstrating improved discrimination abilities under haloperidol. Note that the same pattern emerged when using a continuous variable for WM span, hence for illustrative purposes (Fig. 2) we proceeded with the binary split.

**Figure 2.**
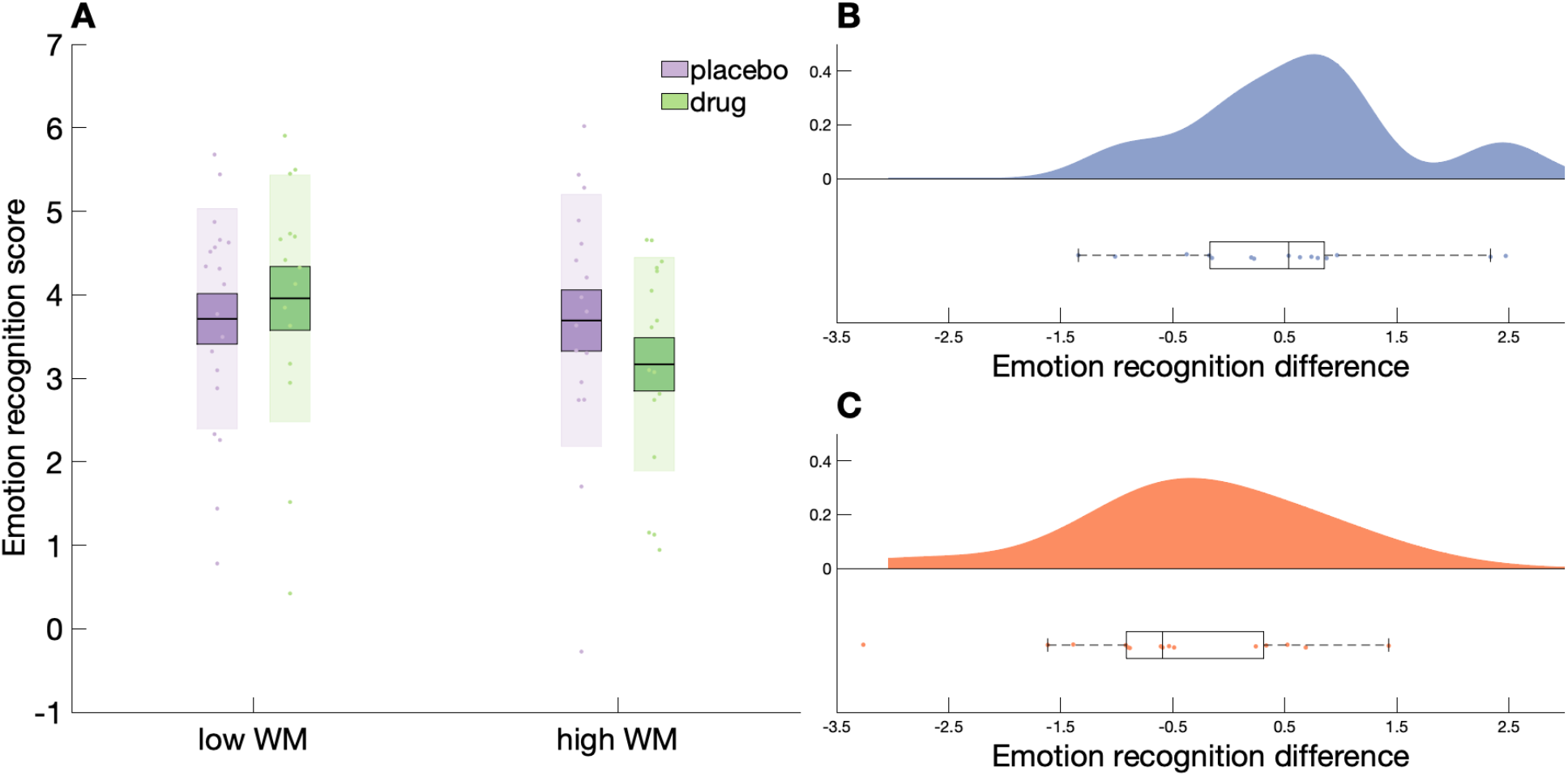
(A) Mean emotion recognition scores for placebo and haloperidol trials by WM group. (B-C) Probability density function (PDF) of emotion recognition difference scores for low (B) and high (C) WM groups. *Note.* (A) Boxes represent 1 SEM above and below the mean (i.e., horizontal lines within boxes), shaded areas surrounding boxes represent 1 SD above and below mean values. (B-C) Emotion recognition difference scores represent the difference between emotion recognition scores in haloperidol and placebo trials, where positive difference scores indicate enhanced emotion recognition performance under haloperidol. The central mark of each of the box plots below PDFs represents the median of each group, edges represent 25^th^ (Q_1_) and 75^th^ (Q_3_) percentiles. Whiskers denote ranges of Q_3_ + 1.5 x (Q_3_ - Q_1_) above and Q_1_ + 1.5 x (Q_3_ - Q_1_) below box edges.

### Effects of haloperidol on movement- and counting-based indices of temporal processing

#### Movements

A Bayesian mixed effects model of drug (PLA, HAL; dummy coded) and WM group (low, high; deviation coded) predicting walking speed indicated a trend towards a negative main effect of drug (*Eµ*_*PLAvsHAL*_ = -0.04, CrI = [-0.08, 0.01], P(*Eµ*_*PLAvsHAL*_) < 0 = 0.94), indicating that, overall, haloperidol tended to reduce walking speed. In addition, there was a trend towards a main effect of WM group (*Eµ*_*WMgroup*_ = 0.08, CrI = [-0.02, 0.19], P(*Eµ*_*WMgroup*_ > 0) = 0.94), demonstrating that under placebo, the low WM group tended to exhibit a slower walking pace relative to high WM individuals (low WM: mean [M] = 1.05 m/s, high WM: M = 1.13 m/s). There further was an interaction between drug and WM group (*Eµ*_*PLAvsHAL,WMgroup*_ = 0.09, CrI = [0.00, 0.18], P(*Eµ*_*PLAvsHAL,WMgroup*_ > 0) = 0.98). Separate post-hoc models for low and high WM groups indicated that, whereas the drug slowed movement speed in the low WM group, there were no drug effects on movement in the high WM group (*Eµ*_*PLAvsHAL,lowWM*_ = -0.08, CrI = [-0.16, -0.01]; *Eµ*_*PLAvsHAL,highWM*_ = 0.01, CrI = [-0.4, 0.6]; Fig. 3A).

**Figure 3.**
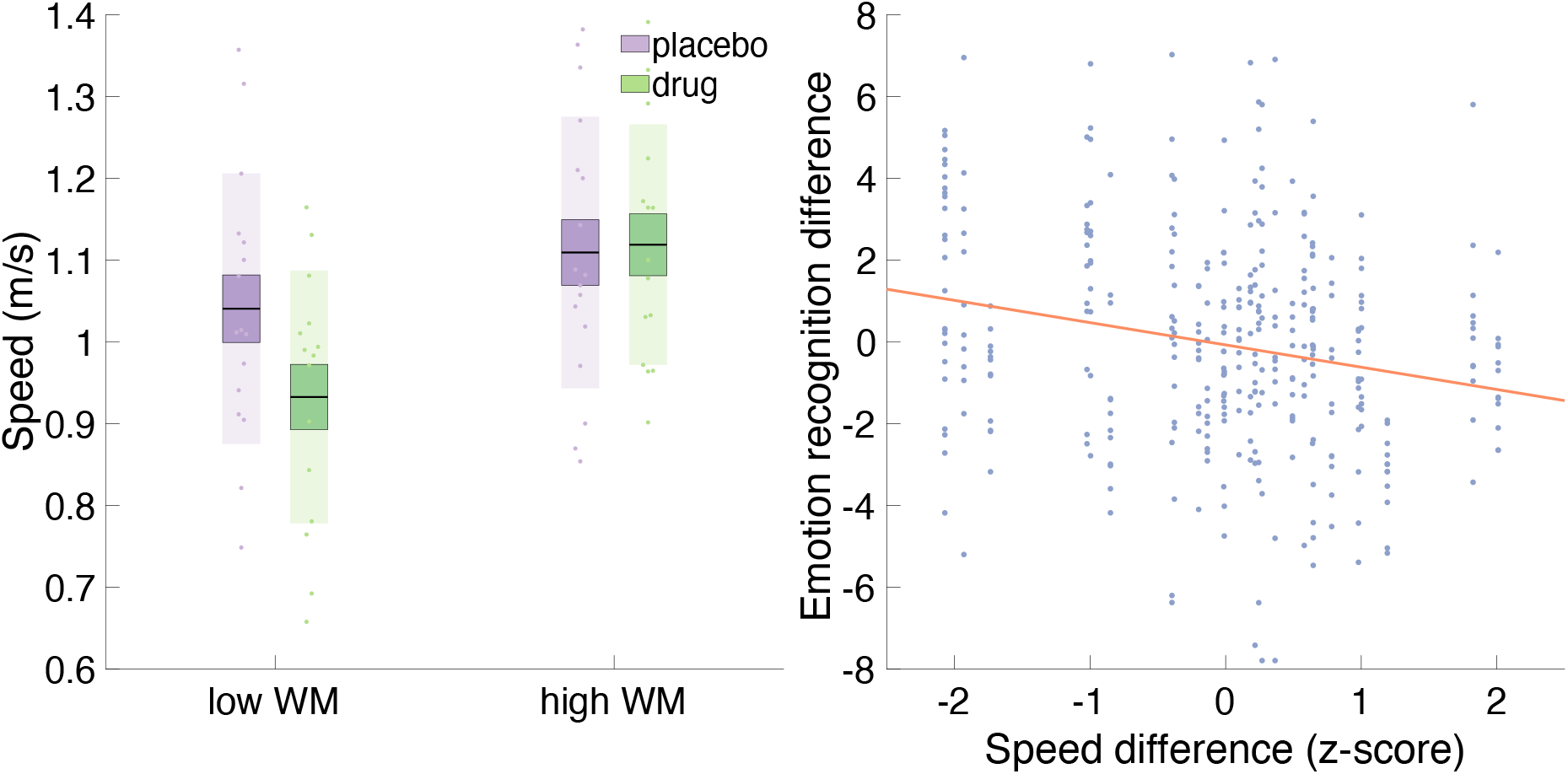
(A) Drug effects on walking speed by WM group. (B) Relationship between drug effects on walking speed and drug effects on emotion recognition scores by WM group. *Note.* (A) Boxes represent 1 SEM above and below the mean (i.e., horizontal lines within boxes), shaded areas surrounding boxes represent 1 SD above and below mean values.

To investigate whether influences of the drug on emotion recognition were modulated by performance on our movement-based index of temporal processing, we first created an index of individual drug effects on emotion recognition by subtracting emotion recognition scores of PLA trials from emotion recognition scores of HAL trials (i.e., *emotion recognition difference scores*). Positive emotion recognition difference scores thus indicate enhanced emotion recognition under haloperidol. Second, we estimated drug effects on walking speed by subtracting mean speed values from HAL trials from mean speed values from PLA trials for each of the two walks (i.e., speed difference values), where negative speed difference values reflect decreased walking speed in HAL relative to PLA trials. Finally, we added speed difference as a covariate to a Bayesian mixed effects model (random effects for subject ID) fitted to emotion and WM group as well as all two- and three-way interactions, predicting emotion recognition difference scores. As noted in the Introduction, previous studies (Edey et al., 2017; Edey et al., 2020) have suggested that emotion recognition relies upon yoking to one’s own movements (whereby slow walkers perceive fast movements as intensely angry). This suggests a possible interaction between emotion and speed difference such that individuals whose movements are slowed under haloperidol exhibit increased emotion recognition difference scores for angry (i.e., fast angry stimuli appear intensely angry), and decreased emotion recognition difference scores for sad PLWs. This exploratory hypothesis was not supported. The first model revealed no interactions with emotion, therefore all following results are based on a model excluding this factor. There was a main effect of WM group, confirming the dependency of drug effects on WM group as reported above (*Eµ*_*WMgroup*_ = -0.79, CrI = [-1.59, -0.00], P(*Eµ*_*WMgroup*_ < 0) = 0.98). Furthermore, there was a main effect of speed difference, indicating that drug effects on walking speed were negatively related to drug effects on emotion recognition (*Eµ*_*speedDiff*_ = -0.66, CrI = [-1.19, -0.12], P(*Eµ*_*speedDiff*_ < 0) = 0.99). A lack of interaction between WM group and speed difference scores (*Eµ*_*speeciDiff,WM group*_ = 0.44, CrI = [-0.30, 1.21]) suggests that the relationship between drug effects on movement and drug effects on emotion perception did not depend on WM span. Thus, in both the high and low WM groups, slower movement speed under the drug was associated with increased emotion recognition, however, haloperidol-induced slowing was observed only in the low WM group (Fig. 3B).

#### Timing

Time perception scores were calculated by subtracting the estimate provided by the participant from the actual duration of a given interval. Time perception difference scores were calculated as the difference between time perception scores in HAL and PLA trials, where negative time perception difference scores reflect slowed time perception under haloperidol. A model with time perception difference scores added as a covariate revealed a marginal negative effect for time perception difference scores for happy PLWs only (*Eµ*_*timeDiff,happy*_ = -0.37, CrI = [-0.73, 0.00]).

## Discussion

The current study tested whether the dopamine D2 receptor antagonist haloperidol modulated emotion recognition from dynamic, whole-body, motion cues. As predicted, the influence of haloperidol on emotion recognition was dependent upon working memory stratification. In our low WM group emotion recognition improved under haloperidol, whereas performance deteriorated in the high WM group. The low WM group also demonstrated slower own movements under the drug, with no impact of haloperidol on walking pace in the high WM group.

To the best of our knowledge our study is the first to illustrate a clear behavioral impact of dopaminergic manipulation on the recognition of numerous emotions and our results thereby highlight the critical importance of accounting for individual differences in measures thought to reflect baseline dopamine function. Such results are consistent with an effect of a dopamine antagonist on emotion recognition previously reported in a sample of 14 males (Lawrence et al., 2002). However, whereas Lawrence et al.’s results were restricted to anger recognition we demonstrate effects across emotions, likely due to accounting for individual differences in baseline dopamine levels. Indeed, our analyses revealed only an interaction between drug and working memory span, and no main effect of drug. Thus, previous mixed neural findings and the absence of behavioral effects likely reflect such individual differences in drug response.

The observation that the low WM group exhibiting an improvement in emotion recognition also slowed their own walking pace is potentially informative with respect to the underlying mechanism. Our results illustrated a negative relationship between drug effects on movement and drug effects on recognition of all three emotions. The crucial role for the motor system in time perception has received widespread recent attention (De Kock et al., 2021), such that a temporal influence of haloperidol on movement performance is likely to reflect wider influences on temporal encoding. Consequently, haloperidol induced movement slowing may signify a slowing of internal timing processes. Furthermore, we observed a similar relationship between individual drug effects on supra-second time perception and drug effects on recognition of happy PLWs, where slower time perception under haloperidol was associated with increased emotion recognition. Thus, effects of haloperidol on both movement- and counting-based timing tasks suggest that slowing of temporal processing is coupled with enhanced emotion recognition. Speculatively, slowed time perception may have led to increased emotion discrimination by enhancing individuals’ sensitivity to temporal cues conveyed in the PLWs.

Notably we did not observe that haloperidol-related movement slowing had emotion-specific effects on recognition (slowing simply predicted improved recognition across *all* emotions). Such emotion-specific effects would have been interesting given our previous work (Edey et al., 2017; Edey et al., 2020) which indicated that we recognize emotions according to comparisons between observed kinematic features and one’s own baseline kinematics – e.g., if the kinematics are faster than the observer’s baseline movement kinematics the model must be angry because this is the speed at which the observer themself feels anger. To be consistent with this, haloperidol-induced slowing should have improved recognition of fast emotions (e.g., anger) yet impaired recognition of slow emotions (e.g., sadness). Our results, however, did not reveal such an interaction between emotion (depicted in the PLW) and drug effects on movement speed. Nevertheless, given the likelihood that one builds models for emotion recognition across a lifetime of experience (Hunnius and Bekkering, 2014; Edey et al., 2020), artificially slowing one’s movement pace in a particular setting (e.g., via haloperidol administration) would be unlikely to re-anchor all models. Given these concerns, we did not feel confident to make strong predictions about emotion-specific effects and we are, indeed, unsurprised to see that this pattern was not reflected in the data.

An important question concerns why we would see such dramatically different results in individuals with high versus low working memory. Interestingly, despite the absence of an effect of haloperidol on movement speed in the high working memory span group, we nevertheless observed that the drug impaired emotion recognition in this group. Thus, suggesting that timing/movement-based effects are not the only mechanism by which haloperidol can affect emotion recognition. One additional mechanism concerns haloperidol’s effects on the maintenance of mental representations. Biologically-inspired models (Frank et al., 2001; Frank, 2005; Frank and Claus, 2006; O’Reilly and Frank, 2006) categorize the effects of haloperidol on mental representations in terms of putative pre-synaptic (i.e., primary blocking of autoreceptors leading to increased dopamine transmission) and post-synaptic (i.e., blocking of post-synaptic heteroreceptors resulting in decreased dopamine transmission) drug effects. Pre-synaptic effects should correspond to enhanced updating of mental representations linked to dopamine bursts (e.g., representations that are rewarded or highly salient: Bromberg-Martin et al. (2010); Diederen and Fletcher (2020)). Post-synaptic effects should result in stable representations that are robust against interference from non-target information. Frank and O’Reilly (Frank and O’Reilly, 2006) have argued that low-span subjects exhibit significantly greater responses to haloperidol (indexed by prolactin, an indirect measure of dopamine levels: Nilsson et al. (1996)) than high-span subjects and that higher doses are more likely to result in both pre- and post-synaptic effects. Since we used a slightly higher dose than Frank and O’Reilly (2.5 mg, versus 2 mg) it is feasible that our low-span subjects obtained a high enough dose of haloperidol that they experienced *both* pre- and post-synaptic effects, whereas our high-span subjects experienced only mild pre-synaptic effects. It would follow from this that our low-span subjects should exhibit enhanced updating of rewarded/salient mental representations (the pre-synaptic effect) *and* more stable representations in general that are robust against interference from non-target information (the post-synaptic effect). In contrast, our high-span participants should have only experienced the former (pre-synaptic) effect.

For accurate emotion recognition, in the context of our paradigm, one must maintain a stable and robust representation of the target PLW (e.g., angry PLW), and resist replacing it with a non-target representation (for example, an imagined PLW prompted by a sad or happy rating scale). Thus, post-synaptic effects, which promote stable and robust mental representations would benefit emotion recognition, resulting in the pattern (high target ratings and low non-target ratings) we observed in our low-span group. In contrast, since pre-synaptic effects favor flexible, rapidly updated, representations they are more likely to result in the pattern we observed in the high-span group where the target and non-target ratings are confused. Consequently, models of the role of dopamine in the updating of mental representations (Frank et al., 2001; Frank, 2005; Frank and Claus, 2006; O’Reilly and Frank, 2006) offer a potential explanation for the differing effects we observe in the high and low-span group, and a potential pathway to explain drug effects on emotion recognition in the absence of effects on temporal processing.

Although the importance of accounting for individual differences in baseline dopamine levels has received widespread attention in other domains of cognition (Williams and Goldman-Rakic, 1995; Kimberg et al., 1997; Mattay et al., 2000; Cools et al., 2008), this study comprises the first illustration within the domain of emotion recognition. We observed that slowed temporal processing under haloperidol was associated with increased emotion recognition, indicating that drug effects on emotion perception could, at least in part, be mediated by effects on movement/timing mechanisms. However, a decline in emotion recognition performance in the absence of drug effects on movement speed in high WM individuals suggests that other mechanisms must also be at play. This work paves the way for future studies to examine how such effects play out with different types of emotion stimuli including static emotion snapshots wherein timing-based mechanisms are less relevant.

## Acknowledgements

This project has received funding from the European Union’s Horizon 2020 Research and Innovation Programme under ERC-2017-STG Grant Agreement No 757583 (Brain2Bee; Jennifer Cook PI).

## Author Contributions

J.L.C., B.S., S.S., and C.P. were substantially involved in the conceptualization of the research idea and design. C.P. and P.H. provided task materials. B.S., S.S., and A.J.R. conducted the investigation process including data acquisition. D.S.F. contributed to the development of the data acquisition methodology and conducted the data processing. B.S. performed the data analysis and prepared the initial publication draft. J.L.C., B.S. and C.P. were involved in revising the work critically, while J.L.C. gave final approval of the version to be published. B.S. agrees to be accountable for all aspects of the work.

## Footnotes

1 https://apps.apple.com/us/app/sensorlog/id388014573

2 The term credible interval comprises the Bayesian analogue of a classical confidence interval, except that probability statements can be made based upon it (see Nalborczyk L, Batailler C, Lœvenbruck H, Vilain A, Bürkner PC (2019) An Introduction to Bayesian Multilevel Models Using brms: A Case Study of Gender Effects on Vowel Variability in Standard Indonesian. J Speech Lang Hear Res 62:1225-1242.)

## Notes

**Conflict of interest:** The authors declare no competing financial interests.

https://osf.io/gcvyj/

### Competing Interest Statement

The authors have declared no competing interest.

